# ENKIE: A package for predicting enzyme kinetic parameter values and their uncertainties

**DOI:** 10.1101/2023.03.08.531697

**Authors:** Mattia G. Gollub, Thierry Backes, Hans-Michael Kaltenbach, Jörg Stelling

## Abstract

Relating metabolite and enzyme abundances to metabolic fluxes requires reaction kinetics, core elements of dynamic and enzyme cost models. However, kinetic parameters have been measured only for a fraction of all known enzymes, and the reliability of the available values is unknown. The ENzyme KInetics Estimator (ENKIE) uses Bayesian Multilevel Models to predict value and uncertainty of *K_M_* and *k_cat_* parameters. Our models use five categorical predictors and achieve prediction performances comparable to deep learning approaches that use sequence and structure information. They provide accurate uncertainty predictions and interpretable insights into the main sources of uncertainty. We expect our tool to simplify the construction of priors for Bayesian kinetic models of metabolism.

## 1 Introduction

To construct dynamic mathematical models for metabolic networks (Kim *et al.*, 2018) or to relate reaction fluxes to metabolite and enzyme abundances in protein cost models (Noor *et al.*, 2016), a main challenge is to find kinetic parameter values for *k_cat_* (rate constants) and *K_M_* (Michaelis constants). Kinetics databases such as BRENDA (Chang *et al.*, 2021) (parameter values for ~ 90000 enzymes from > 150’000 publications) and SABIO-RK (Wittig *et al.*, 2018) (~ 7500 publications) cover only a minority of known enzymes (e.g., in few species).

To predict parameter values from existing databases, a recent combination of machine- and deeplearning models achieved an coefficient of determination (R^2^) of 0.53 for affinities (Kroll *et al.*, 2021) and of 0.40 for catalytic rates (Kroll *et al.*, 2022) on independent test sets. Unfortunately, R^2^ (or Root Mean Square Error (RMSE)) values do not reflect how well the underlying data determines individual parameter values. To assess prediction uncertainties of kinetic models, one must account for uncertainties due to (unknown) annotation inaccuracies, experimental errors, and chemical environments.

Here, we develop Bayesian Multilevel Models (BMMs) that capture a simple hierarchy of enzyme properties. They predict *K_M_* and *k_cat_* values as well as their uncertainties through the linear combination of *population-level* effects (with a single value for the entire population) and *group-level* effects (with different values across groups and a common normal distribution). The models can be easily used through ENKIE, a python package for the estimation of thermodynamically consistent enzyme kinetic parameter values and their uncertainties.

## 2 Methods

### Statistical models

We use hierarchical BMMs to express the relationship between the properties of a kinetic parameter, *K_M_* or *k_cat_*, and its value (Supplementary Methods). The models capture the reduction in variability as the classification of a parameter value becomes more precise, without considering structural and physical properties of enzymes, reactions, and metabolites explicitly. For *K_M_* and *k_cat_* values, we assume normally distributed residuals with mean *μ*_*_ and standard deviation *σ*_*_ (* ∈ {*M, cat*}). We describe *μ*_*_ by a population average and a series of nested group-level effects. For *K_M_*, the substrate is the first grouping level because parameter values for a substrate are conserved across reactions (Park *et al.*, 2016). We then nest the Enzyme Commission (EC)-reaction pair, the protein family, and the protein identifier denoting a specific enzyme of a specific organism. For *k_cat_*, we replace the substrate with a nested hierarchy of the first three components of the EC number and include the reaction direction to distinguish between forward and backward rates. To model *σ*_*_, we use two different population averages for entries with / without an annotated protein identifier because entries with unknown identifier are presumably harder to fit. We use a group-level effect to incorporate that the size of the residuals can differ across reactions because residuals capture many unmodeled factors, such as experimental errors, different experimental conditions, and annotation errors.

### Fitting and evaluation

The BMMs capture model uncertainty (confidence in the estimated model parameters given the data) and the residuals *σ*_*_ (discrepancy between measurements of the same parameter, possibly because of unmodeled effects). Because these can be strongly correlated, we fitted the models in a Bayesian framework. Specifically, we implemented the models in R and fitted them using the Markov Chain Monte Carlo (MCMC)-based `brms` package (Bürkner, 2017) to data extracted from BRENDA and SABIO-RK (Supplementary Methods). The simulated chains had no divergent transitions and a potential scale reduction factor < 1.05, suggesting good fit quality (Gelman *et al.*, 2013). For model analysis, we omitted parameter balancing to simplify interpretation and benchmarking (see Supplementary Methods).

### Implementation

ENKIE (Fig. 1a) uses generally available inputs: reaction stoichiometries, metabolite / reaction identifiers in any namespace supported by MetaNetX (Moretti *et al.*, 2021), ECs, protein identifiers and, optionally, physiological properties of the reaction compartments (pH, pMg, temperature and ionic strength). It uses MetaNetX to standardize identifiers and Uniprot (The UniProt Consortium, 2021) to obtain protein family annotations. Inputs are passed to the `brms predict` method via the `rpy2` package. Optionally, samples from the joint distribution of predicted parameters are combined with free energy estimates from eQuilibrator (Beber *et al.*, 2022) using parameter balancing (Lubitz *et al.*, 2010) to obtain a multivariate normal distribution of thermodynamically consistent estimates and uncertainties of free energies and kinetic parameters.

**Figure 1.**
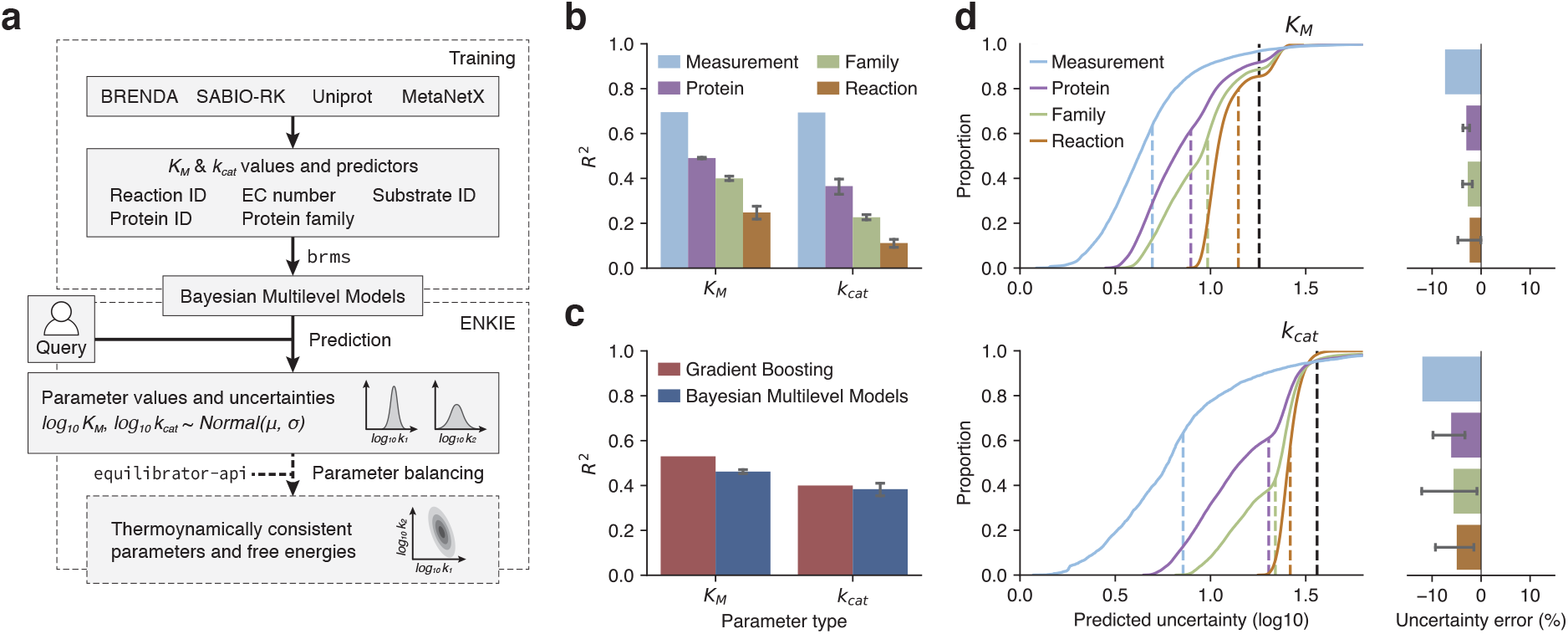
ENKIE design and performance. **a**, Overview. We gather kinetic parameter values from BRENDA and SABIO-RK, use Uniprot and MetaNetX to consistently annotate them, and train models for value and uncertainty of *K_M_* and *k_cat_* parameters. The `enkie` package uses the models to predict value and uncertainty for query parameters, optionally with eQuilibrator (Beber *et al.*, 2022) and parameter balancing (Lubitz *et al.*, 2010) to construct thermodynamically consistent parameters. **b**, *R*^2^ for ENKIE when predicting parameter values for measurements, proteins, protein families, and reactions not present in the training set. **c**, *R*^2^ for methods based on gradient boosting (Kroll *et al.*, 2021, 2022) and our BMMs. **d**, Empirical cumulative distribution of the predicted parameter uncertainties on different dataset split strategies (left). Dashed lines: standard deviation of parameter values over the entire dataset (black) and RMSE (colored). Right, error in the predicted uncertainties (deviation from one in the Root Mean Square Normalized Error (RMSNE)).

## 3 Results

We first analyzed the features’ contributions to the predicted *K_M_* and *k_cat_* values (Table S1). All population- and group-level effects were estimated with small uncertainty. The strongest determinants (i.e. the group-level effects with the largest average size) for *K_M_* and *k_cat_* were the substrate and reaction identifier respectively. Importantly, the standard deviations of the only organism specific group-level effects (the Uniprot identifier, Table S1) were significantly smaller than the standard deviations over the entire dataset (Table S5). Hence, measurements from characterized organisms are informative for predicting values in new organisms.

In terms of training data quality, the average values for the estimated residuals *σ_M_* and *σ_cat_* were slightly larger with missing protein identifier annotations (Table S1), justifying the split into two groups. For entries with a protein identifier (which we expect to be always available in practical applications), estimated upper bounds of the 95% Confidence Intervals (CIs) above one imply that, for certain parameters, multiple measurements with the same protein span several orders of magnitude.

Next, we characterized the performance of our BMMs at predicting values for completely unknown measurements, proteins, protein families, and reactions. Specifically, we performed 5-fold cross-validation with different types of *folds* (equally sized partitions of the input dataset; Supplementary Methods and Fig. S1). Fig. 1b shows that the models remain predictive even when extrapolating to new reactions. Grouped folds are notoriously challenging and a drop in performance is expected. *K_M_* models generally achieve a better *R*^2^ than *k_cat_* models. This could be because affinities are more conserved across organisms than rates, as shown by the lower variance explained by protein effects in *K_M_* models (13.2%) than in *k_cat_* models (23.9%) (Table S1).

We compared the R^2^ achieved by BMMs and state of the art methods (Kroll *et al.*, 2021, 2022) for prediction of *K_M_* and *k_cat_* using the same training and test data reported in the respective publications. As our models only use ECs, identifiers, and protein families, we expect them to have lower performance than methods that leverage sequence and structure information. However, the difference was only marginal for *K_M_*, and surprisingly not significant for *k_cat_* values (Fig. 1c). This suggests that detailed sequence or structure similarities are informative for affinities, but are not more informative than discrete classifications for *k_cat_*.

To evaluate the predicted uncertainties, which are unique to our approach, we first investigated their distribution (Fig. 1d). As expected, the distributions of predicted uncertainties for the folding strategies are approximately centered around the RMSE, but they additionally provide detailed per-prediction uncertainties that can be up to three times lower than the RMSE. We assessed the quality of the predicted uncertainties through use the Root Mean Square Normalized Error (RMSNE) (Supplementary Methods), a value that is expected to be one for perfect predictions. For both *K_M_* and *k_cat_,* in all fold strategies, the RMSNE was close to one (Table S3), and only underestimated the uncertainty by at most 12%(Fig. 1d). This is likely caused by violations in the normality assumptions, for example, because of outliers (misannotated entries) in the databases.

Lastly, we examined the group-level effects of the residuals to determine if there are systematic differences across reactions and organisms, hypothetically due to difficulties in quantifying specific compounds, the speed of reactions, differences across kingdoms, or annotation inaccuracies. We used the already fitted models and additionally fitted models for *K_M_* and *k_cat_* where the group-level effects of the residuals were based on the organism (Supplementary Methods). Fig. S2 shows clear differences among reaction groups. For example, pyruvate kinase (MNXR103371) had a large positive effect for the residuals of *k_cat_*, caused by a single publication contributing 52 measurements for the same protein with strong variations in the concentrations of several ions and sugar phosphates. In contrast, the RuBisCO carboxylase and oxigenase reactions (MNXR191185 and MNXR191183) had negative effects for both *K_M_* and *k_cat_*, suggesting that the enzyme has been characterized with particular precision. For organism effects, we did not observe systematic difference across kingdoms (Fig. S2).

## 4 Discussion

Our approach builds on linear regression to predict *K_M_* values (Borger *et al.*, 2006), but introduces group-level effects. This enables prediction qualities comparable to current deep-learning methods that additionally use enzyme sequence and structure information. Importantly, our method provides reliable uncertainty estimates; those uncertainties can be up to three times smaller than the RMSE on the log_10_ scale.

Compared to machine learning approaches, our models are easily interpretable. The effect sizes of the fitted models directly inform us about how much variance is explained by simple features such as substrate, reaction and protein family. We also showed how the effects on the residuals can be directly explained with the data. This is important because, in addition to ambiguously defined parameters, we frequently encountered challenges in the standardization and interpretation of database values. The residuals estimated by our Bayesian Multilevel Models (BMMs) highlight inconsistencies and can thus help improve the structure and accuracy of enzyme databases.

ENKIE does not cover activation and inhibition constants, but we expect that the approach can be extended. Like all methods trained on measurable values, it likely yields biased predictions for extreme values (e.g., backward *k_cat_* of highly favourable reactions). A possible solution could include standard reaction energies as predictors. Even with these limitations, ENKIE estimates can be used directly to increase the robustness of applications that rely on kinetic parameters.

## Availability

Code and Python package are available at gitlab.com/csb.ethz/enkie and pypi.org/project/enkie.

## Acknowledgements

We thank Janina Linnik, Lukas Widmer, and Sebastian Weber for helpful discussions.

## Funding

This work was supported by the Swiss National Science Foundation Sinergia project #177164.

## Notes

https://gitlab.com/csb.ethz/enkie

### Competing Interest Statement

The authors have declared no competing interest.

