## Supplementary Information for "ENKIE: A package for predicting enzyme kinetic parameter values and their uncertainties"

Mattia G. Gollub [1], Thierry Backes [1], Hans-Michael Kaltenbach [1] and Jörg Stelling [1,*]

[1]Department of Biosystems Science and Engineering and SIB Swiss Institute of Bioinformatics, ETH Zurich, Basel, 4058, Switzerland.

#### Supplementary Methods

##### Statistical models

Using the brms notation for Multilevel Models (MLMs) we can write the models for the $\log_{10}$ values of $K_M$ and $k_{cat}$ as

$$\begin{aligned}
\log_{10}(K_M) &\sim \mathcal{N}(\mu_M, \sigma_M) \\
\mu_M &\sim 1 \\
&\quad + (1|\texttt{substrate}) \\
&\quad + (1|\texttt{substrate}:\texttt{ec}:\texttt{reaction}) \\
&\quad + (1|\texttt{substrate}:\texttt{ec}:\texttt{reaction}:\texttt{family}) \\
&\quad + (1|\texttt{substrate}:\texttt{ec}:\texttt{reaction}:\texttt{family}:\texttt{uniprot_ac}) \\
\log(\sigma_M) &\sim 0 + \texttt{has_ac} + (1|\texttt{substrate}:\texttt{ec}:\texttt{reaction})
\end{aligned} \tag{1}$$

$$\begin{aligned}
\log_{10}(k_{cat}) &\sim \mathcal{N}(\mu_{cat}, \sigma_{cat}) \\
\mu_{cat} &\sim 1 \\
&\quad + (1|\texttt{ec1}) \\
&\quad + (1|\texttt{ec1}:\texttt{ec2}) \\
&\quad + (1|\texttt{ec1}:\texttt{ec2}:\texttt{ec3}) \\
&\quad + (1|\texttt{ec}:\texttt{reaction}:\texttt{is_forward}) \\
&\quad + (1|\texttt{ec}:\texttt{reaction}:\texttt{is_forward}:\texttt{family}) \\
&\quad + (1|\texttt{ec}:\texttt{reaction}:\texttt{is_forward}:\texttt{family}:\texttt{uniprot_ac}) \\
\log(\sigma_{cat}) &\sim 0 + \texttt{has_ac} + (1|\texttt{ec}:\texttt{reaction}:\texttt{is_forward})\,.
\end{aligned} \tag{2}$$

Both $\log_{10}(K_M)$ and $\log_{10}(k_{cat})$ values are assumed to have normally distributed residuals with standard deviations $\sigma_*$ around their mean values $\mu_*$ ($* \in \{M, cat\}$). Mean values are described by a population average ("`1`") and a series of nested group-level effects ("$+$(`1`|...)"). For $K_M$, we first assume that parameter values for a substrate are generally conserved across species (Park *et al.*, 2016) and thus use the substrate as the first level of grouping. We then expand the hierarchy by further nesting the Enzyme Commission (EC)-reaction pair, the protein family, and lastly the protein identifier denoting a specific enzyme of a specific organism. We use a similar model for $k_{cat}$, with the difference that we replace the substrate effect at the root with a nested hierarchy of the first three components (`ec1`, `ec2`, and `ec3`) of the EC number. Additionally we include the reaction direction to distinguish between forward and backward rates.

Similarly, we model $\sigma_*$ using both population- and group-level effects. We use two different population averages depending on whether reported values have an annotated protein identifier or not ("`0`$+$`has_ac`") to capture the assumption that entries with unknown identifier are harder to fit. Furthermore, we use a group-level effect to incorporate the assumption that the size of the residuals can differ across reactions. Residuals subsume many unmodeled factors, such as experimental error, differences in experimental conditions, or annotation errors. Each of those can be reaction dependent. For example, some reactions may require different measurement techniques, be more or less sensitive to pH and temperature variations, or be more prone to reporting errors when they are catalyzed by different isozymes. As these differences are unknown *a priori*, we estimate them from the data.

The models Eq. (1) and (2) contain two sources of uncertainty: the uncertainty caused by the residuals (quantified by $\sigma_*$) and the model uncertainty. The former captures the residual variance of the unmodeled effects, while the latter describes how confident we are in the model parameters given the data we have. These uncertainties can present strong correlations, for example, when parameters are not identifiable, which are hard to estimate explicitly with maximum likelihood methods (Bates *et al.*, 2015). Therefore, we fit the models in a Bayesian framework, which allows us to sample model parameters from their posterior distribution. As a consequence, during prediction, the models can seamlessly estimate values for measured and unknown parameters while providing uncertainties for the estimates. For example, predicting a value for which measurements are available consists in summing effects samples to obtain samples of $\mu_*$ and $\sigma_*$, then sampling from $\mathcal{N}(\mu_*, \sigma_*)$. We then compute the final predicted values and uncertainties $\hat{y}_*$ and $\hat{\eta}_*$ as the mean and standard deviation of the samples. If one or more group-level effects are unknown, for example, because a specific protein has no measurement, then the procedure is the same except that the missing effects are sampled from their distribution.

#### Data collection and processing

We collected the values and features used for training from SABIO-RK (Wittig *et al.*, 2018) (v2.19) and BRENDA (Chang *et al.*, 2021) on January $4^{th}$ 2023. The features and main statistics of the final dataset are described in Table S4 and Table S5, respectively.

For SABIO-RK, we downloaded all entries through the SABIO-RK REST API and only kept measurements of $K_M$ and $k_{cat}$ parameters. We converted all metabolite and reaction identifiers to the MetaNetX namespace (Moretti *et al.*, 2021). As the reaction mappings do not necessarily preserve direction, we mapped and compared substrates and products of the SABIO-RK and MetaNetX reactions to identify the correct direction of the reported $k_{cat}$. We then used the `ete3` toolkit (Huerta-Cepas *et al.*, 2016) to translate textual organism annotations to NCBI taxonomy identifiers.

For BRENDA, we retrieved all $K_M$ and $k_{cat}$ entries using the `brendapy` package and used MetaNetX to convert substrate names to MetaNetX identifiers. The mapping was not unique, mainly because of ambiguous names and unspecific isomers. For such cases, we created a copy of the entry for each isomer. The Human Metabolome Database (HMDB) (Wishart *et al.*, 2022), one of the sources used by MetaNetX, contained incorrect names (e.g. MNXM8 is described both as NAD+ and NADH) and its entries were thus excluded from the mapping. As BRENDA does not specify reactions for individual measurements, we inferred reaction identifiers and directions from the EC numbers and the substrate identifiers. We used KEGG (Kanehisa *et al.*, 2021) to map EC numbers to reaction identifiers, as MetaNetX reported several incorrect mappings (e.g. MNXR95726, a NADP-specific dehydrogenase, includes EC 1.1.1.1 which is NAD-specific). Entries with ambiguous reaction assignments were discarded.

In post-processing, we standardized and merged the entries collected from both databases in a single dataset and only retained measurements for wild type proteins (i.e. entries not containing the word ‘mutant’ in the comment or variant fields). As protein identifiers were missing for a significant portion of the dataset, we used the Uniprot search API (The UniProt Consortium, 2021) to infer candidate identifiers from EC and taxonomy identifiers. We then used the Uniprot Retrieve/ID mapping API to obtain protein family annotations for each entry with a protein identifier. Finally, we addressed cases in which the process above did not yield unique results. Where protein identifier were ambiguous, we replaced them with new pseudo-identifiers that were unique for each organism-tissue pair. If protein family information were ambiguous, we removed them. In a last cleanup step, we standardized units, maintained only entries with $K_M$ expressed in molar concentrations or $k_{cat}$ expressed in $s^{-1}$, removed duplicate entries, and converted all values to the $\log_{10}$ scale as kinetic parameters are positive values spanning several orders of magnitude.

Note that our data collection and post-processing workflow expands on previous work (Borger *et al.*, 2006; Heckmann *et al.*, 2018; Kroll *et al.*, 2021; Li *et al.*, 2022) in that neither reported the usage of reaction identifiers (which are not equivalent to EC-substrate pairs[1]), nor inferred protein identifiers and protein families from EC numbers and organism identifiers where possible.

[1]For example, EC 1.1.1.1 (alcohol dehydrogenase) acts on a wide range of alcohols, and its catalytic rate for NAD/NADH depends on the co-substrate and thus on the reaction. EC numbers can also imply additional information on the protein which are not accounted for in the reaction definition. For example, EC 5.4.2.11 and 5.4.2.12 catalyze the same reaction but differ in the presence of a cofactor. However, we include this information at the protein family level.

#### Model fitting

We implemented and fitted the models in `R` with the `brms` package (Bürkner, 2017), a Bayesian framework for fitting different kinds of regression models that uses `stan`'s No-U-Turn Sampler (NUTS) sampler (Hoffman and Gelman, 2011) for estimating model parameters. We used the recommended default settings for sampling (4 chains, 2000 steps, the initial 1000 steps are considered warm-up), only increasing the target average acceptance probability `adapt_delta` to 0.98 to avoid divergences caused by the funnel geometry of the target distribution (Betancourt and Girolami, 2013). Thinning was adjusted to obtain a total of 1000 samples.

Table S6 summarizes the priors used for fitting. The prior means for the population level effects were constructed from the means and average per-parameter standard deviations in the $K_M$ and $k_{cat}$ data. Standard deviations were conservatively set to 0.5. `has_ac` had the same prior for the `True` and `False` cases, so that potential differences between the two classes would emerge from the data. For the priors on the standard deviations of the group-level effects, we set a smaller standard deviation than expected to impose a small degree of regularization, such that group-level effects have a significant size only if supported by the data.

#### Model evaluation

We compared the coefficient of determination ($R^2$) achieved by our models and by state of the art methods (Kroll *et al.*, 2021, 2022) using the training and test datasets reported in the respective publications. In both cases, the authors used a 80% - 20% split. To quantify the uncertainty in the estimated $R^2$, we generated 4 additional splits with the same data (such that there is no overlap between the test datasets of the splits) and reported the 95% Confidence Interval (CI) of the $R^2$ over the 5 splits. We additionally summarized the performances reported by previous works Table S2. However, one must be careful comparing the results, as each publication used different datasets and split strategies.

Furthermore, we characterized the performance of our models at predicting values for completely unknown proteins, protein families and reactions by performing cross-validation with different fold selection strategies. Figure S1 compares predicted and experimental values in the training set and during cross-validation with different fold types. Here, it is important to distinguish between the random folds and the notoriously more challenging grouped folds. In grouped folds, the values of the unknown group-level are sampled from their distribution and have mean zero. Thus, the average of the predicted values is given by the average value for the known groups. This strategy leads to patterns that can be observed in Figure S1. While in the training set we observe the highest density of points on the diagonals, structured folds lead to the emergence of horizontal stripes where all the predictions of unknown groups concentrate.

The Root Mean Square Normalized Error (RMSNE) to evaluate the quality of the predicted uncertainties is defined as the Root Mean Square Error (RMSE) computed on residuals that have been divided by the prediction uncertainty:

$$\text{RMSNE} = \sqrt{\sum_i \delta_i^2}, \qquad \delta_i = \frac{\hat{y}_i - y_i}{\hat{\eta}_i} \,, \tag{3}$$

where $y_i$, $\hat{y}_i$, $\hat{\eta}_i$ and $\delta_i$ are the measured value, the predicted value, the predicted uncertainty and the normalized residual of entry $i$. With perfect uncertainty predictions, this value should be equal to one, while smaller or larger values indicate over- and under-estimation of the uncertainty. The error in the predicted uncertainties was then computed as $error = (1 - \text{RMSNE}) \cdot 100$, such that positive and negative values indicate overestimation and underestimation of the uncertainty.

To examine the group-level effects of the residuals, we used the already fitted models and additionally fitted models for $K_M$ and $k_{cat}$ where the group-level effects of the residuals were based on the organism:

$$\begin{aligned} \log(\sigma_M) &\sim 0 + \texttt{has\_ac} + (1|\texttt{organism}) \\ \log(\sigma_{cat}) &\sim 0 + \texttt{has\_ac} + (1|\texttt{organism}) \,. \end{aligned} \tag{4}$$

For both types of models, we selected the 10 groups with the highest number of observations to minimize noise.

### Supplementary Figures

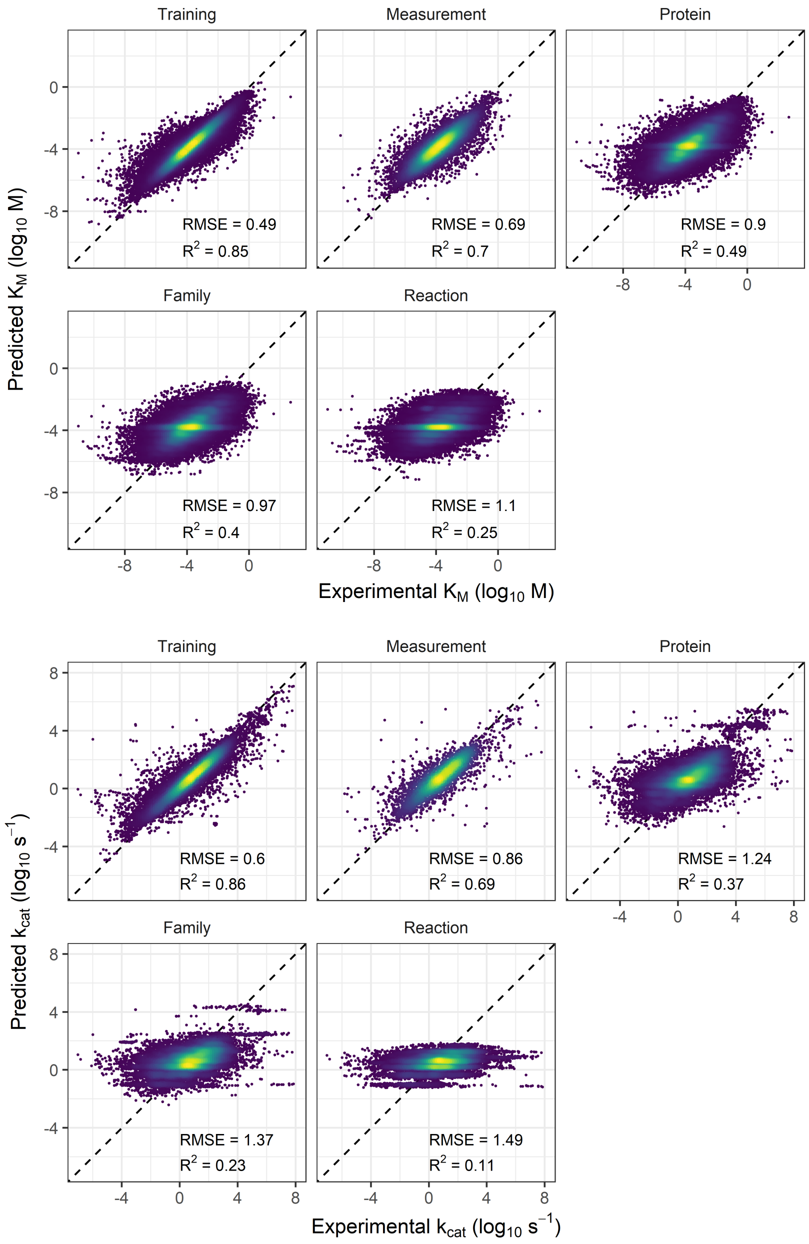


**Figure S1.** Comparisons of predicted and experimental values for $K_M$ (top) and $k_{cat}$ (bottom). The plots show the performance of the model on the training set and during cross-validations with folds selected at random or systematically partitioning the data by protein identifier, protein family and reaction. Lighter colors denote higher points density.

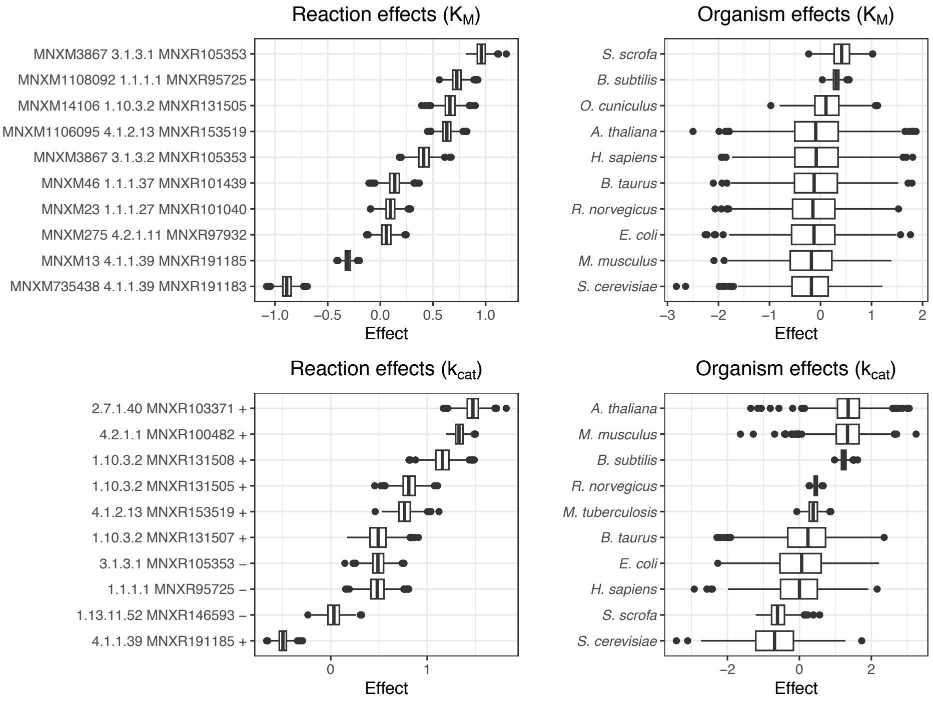


**Figure S2.** Effect on the residuals of $K_M$ (top) and $k_{cat}$ (bottom) models using reactions (left) or organisms (right) as predictors for the ten groups with the highest number of measurements. Reaction groups are defined by the triples metabolite - EC - reaction for $K_M$ and EC - reaction - direction (+: forward, - backward) for $k_{cat}$.

#### Supplementary Tables

**Table S1.** Summary of the estimated effects. For group-level effects ((1|...)) we report the standard deviation of the effect distribution, which is also the root-mean-square effect size. For the population-level effects we report the estimated effect. Individual group-level effects are not reported. The Error column reports the uncertainty in the estimated effect. The last column reports the percentage of variance explained by each group-level effect (excluding the residuals), computed as the variance of an effect divided by the total variance of all effects.

| Effect | | Value | Error | Variance explained |
|---|---|---|---|---|
| $\mu_M$ | 1 | -3.83 | 0.02 | - |
| $\mu_M$ | (1\|substrate) | 0.92 | 0.01 | 57.9% |
| $\mu_M$ | (1\|substrate : ec : reaction) | 0.60 | 0.01 | 24.6% |
| $\mu_M$ | (1\|substrate : ec : reaction : family) | 0.25 | 0.01 | 4.3% |
| $\mu_M$ | (1\|substrate : ec : reaction : family : uniprot_ac) | 0.44 | 0.00 | 13.2% |
| $\log(\sigma_M)$ | has_ac = False | -0.85 | 0.01 | - |
| $\log(\sigma_M)$ | has_ac = True | -0.99 | 0.01 | - |
| $\log(\sigma_M)$ | (1\|substrate : ec : reaction) | 0.64 | 0.01 | - |
| $\mu_{cat}$ | 1 | 0.51 | 0.10 | - |
| $\mu_{cat}$ | (1\|ec1) | 0.19 | 0.06 | 2.0% |
| $\mu_{cat}$ | (1\|ec1 : ec2) | 0.23 | 0.06 | 3.0% |
| $\mu_{cat}$ | (1\|ec1 : ec2 : e3) | 0.50 | 0.04 | 13.7% |
| $\mu_{cat}$ | (1\|ec : reaction : is_forward) | 0.91 | 0.02 | 45.4% |
| $\mu_{cat}$ | (1\|ec : reaction : is_forward : family) | 0.47 | 0.02 | 12.1% |
| $\mu_{cat}$ | (1\|ec : reaction : is_forward : family : uniprot_ac) | 0.66 | 0.01 | 23.9% |
| $\log(\sigma_{cat})$ | has_ac = False | -0.83 | 0.03 | - |
| $\log(\sigma_{cat})$ | has_ac = True | -1.02 | 0.02 | - |
| $\log(\sigma_{cat})$ | (1\|ec : reaction : is_forward) | 0.79 | 0.02 | - |

**Table S2.** Summary of the performances reported by published methods and of the models for $K_M$ and $k_{cat}$ developed in this work. All publications used different train/test datasets. Where reported by the authors, performances are measured through the coefficient of determination ($R^2$) (higher is better) and the Root Mean Square Error (RMSE) (lower is better). Where needed, these scores were computed from the reported Pearson correlation coefficients or Mean Square Errors (MSEs). The folds column specifies which kind of grouping was applied to split training and test datasets. k-f: k-fold cross-validation. LOO: Leave-one-out cross-validation. *: These publications used mean/maximum values for validation and/or removed outliers. Therefore, their datasets have smaller variance than the values reported in databases and the RMSE values are expected to be lower.

| | Method | Reference | Folds | $R^2$ | RMSE |
|---|---|---|---|---|---|
| $K_M$ | Multilevel model | This work | Protein (5-f) | 0.49 | 0.90 |
| | Multilevel model | This work | Measurement (5-f) | 0.70 | 0.69 |
| | Gradient boosting* | Kroll *et al.* (2021) | Protein (5-f) | 0.53 | 0.81 |
| | Linear regression | Borger *et al.* (2006) | Random (LOO) | – | 1.01 |
| $k_{cat}$ | Multilevel model | This work | Protein (5-f) | 0.37 | 1.24 |
| | Gradient boosting* | Kroll *et al.* (2022) | Protein (5-f) | 0.4 | 0.92 |
| | Deep learning* | Li *et al.* (2022) | Protein (10-f) | 0.42 | – |
| | Random forest | Heckmann *et al.* (2018) | Protein (10-f) | 0.31 | – |

**Table S3.** Root Mean Square Normalized Error (RMSNE) achieved by the models during cross-validation.

| Folds type | $K_M$ | $k_{cat}$ |
|---|---|---|
| Measurement | 1.07 | 1.12 |
| Protein | 1.03 | 1.06 |
| Family | 1.03 | 1.06 |
| Reaction | 1.02 | 1.05 |

**Table S4.** Features used for training and prediction in ENKIE.

| Parameter | Type | Description |
|---|---|---|
| `ec` | string | EC number of the enzyme |
| `ec1` | string | First component of EC number ($k_{cat}$ only) |
| `ec2` | string | Second component of EC number ($k_{cat}$ only) |
| `ec3` | string | Third component of EC number ($k_{cat}$ only) |
| `reaction` | string | MetaNetX (Moretti *et al.*, 2021) identifier of the reaction |
| `is_forward` | boolean | `True` if $k_{cat}$ refers to the forward direction, `False` otherwise ($k_{cat}$ only) |
| `substrate` | string | MetaNetX identifier of the substrate ($K_M$ only) |
| `family` | string | Protein family according to Uniprot classification |
| `uniprot_ac` | string | Uniprot accession identifier(s) of the protein(s) |
| `has_ac` | boolean | `False` for entries that do not contain Uniprot identifiers |
| `type` | {`km`, `kcat`} | Parameter type |
| `value` | float | Parameter value, in $\log_{10}$ scale |

**Table S5.** Main properties of the training dataset constructed form SABIO-RK and BRENDA. Statistics are based on molar concentrations ($K_M$) and $s^{-1}$ ($k_{cat}$). The number of BRENDA entries is computed after removing entries that were already present in SABIO-RK.

| | $K_M$ | $k_{cat}$ | Total |
|---|---|---|---|
| SABIO-RK entries | 35 167 | 14 544 | 49 711 |
| BRENDA entries | 34 153 | 11 104 | 45 257 |
| Total entries | 69 320 | 25 648 | 94 968 |
| Average value ($\log_{10}$) | −3.80 | 0.73 | – |
| Standard deviation ($\log_{10}$) | 1.26 | 1.56 | – |
| Number of EC codes | 3 640 | 2 386 | 3 674 |
| Number of reactions | 9 598 | 5 982 | 9 769 |
| Number of substrates | 5 826 | – | – |
| Number of protein families | 1 967 | 1 323 | 1 988 |
| Number of organisms | 2 849 | 1 621 | 2 888 |
| Number of proteins | 12 495 | 6 455 | 12 723 |

**Table S6.** Priors used for fitting the models. *sd* denotes priors on the standard deviation of group-level effects distributions. $\mathcal{N}$ and $\mathcal{N}^+$ are the normal and half-normal distributions.

| Effect | $K_M$ | $k_{cat}$ |
|---|---|---|
| `1` ($\mu_*$) | $\mathcal{N}(-3.8, 0.5)$ | $\mathcal{N}(0.7, 0.5)$ |
| `has_ac` ($\log(\sigma_*)$) | $\mathcal{N}(\log(0.1), 0.5)$ | $\mathcal{N}(\log(0.1), 0.5)$ |
| Group-level sd ($\mu_*$) | $\mathcal{N}^+(0, 0.1)$ | $\mathcal{N}^+(0, 0.1)$ |
| Group-level sd ($\log(\sigma_*)$) | $\mathcal{N}^+(0, 0.1)$ | $\mathcal{N}^+(0, 0.1)$ |
